# Identification and Functional Characterization of lncRNAs involved in Human Monocyte-to-Macrophage Differentiation

**DOI:** 10.1101/2024.06.20.599925

**Authors:** Christy Montano, Sergio Covarrubias, Eric Malekos, Sol Katzman, Susan Carpenter

**Affiliations:** Department of Molecular, Cell and Developmental Biology, University of California Santa Cruz, California, USA; Department of Biomolecular Engineering, University of California Santa Cruz, California, USA

## Abstract

Long noncoding RNAs (lncRNAs) make up the largest portion of RNA produced from the human genome, but only a small fraction have any ascribed functions. Although the role of protein-coding genes in macrophage biology has been studied extensively, our understanding of the role played by lncRNAs in this context is still in its early stages. There are over 20,000 lncRNAs in the human genome therefore, attempting to select a lncRNA to characterize functionally can be a challenge. Here we describe two approaches to identify and functionally characterize lncRNAs involved in monocyte-to-macrophage differentiation. The first involves the use of RNA-seq to infer possible functions and the second involves a high throughput functional screen. We examine the advantages and disadvantages of these methodologies and the pipelines for validation that assist in determining functional lncRNAs.

## Introduction

The advent of RNA-sequencing (RNA-seq) has revealed that long noncoding RNAs (lncRNAs) make up the largest portion of the human transcriptome. According to GENCODE V46, 20,310 lncRNAs are encoded in the human genome. LncRNAs are transcripts greater than 500 nucleotides long, typically lacking protein-coding potential, and are often spliced and polyadenylated (1). LncRNAs are incredibly cell-type specific and have been shown to regulate biological processes, including immunity and cell differentiation, through various mechanisms (2, 3, 4). There are different categories of lncRNAs, including intergenic, antisense, and bidirectional, defined in relation to nearby protein-coding genes. While categorization can offer preliminary clues about a lncRNA gene’s functions, recent research suggests that lncRNA loci are multifaceted with functions emerging from the RNA itself, the act of transcription, through enhancer related mechanisms, and in some cases, even by encoding small peptides (5, 6, 7, 8). Therefore, it is vital to establish rational guidelines and effective methodologies for exploring the functional roles of lncRNAs.

Monocytes and macrophages play pivotal roles in the innate immune response, acting as initial responders to foreign pathogens (9). The process of monocyte-to-macrophage differentiation requires precise regulation. A dysregulated differentiation response can trigger an exaggerated inflammatory response and blood cancers like leukemia (10, 11). While previous research has focused on the contribution of protein-coding genes in monocyte differentiation, we sought to contribute to early work describing lncRNA contributions to this process (12).

We describe two pipelines to identify functionally relevant lncRNAs and outline what we believe are the most important criteria to consider when choosing which lncRNAs to pursue further. We utilized THP1 cells, a human acute monocytic leukemia cell line, to identify lncRNA regulators of monocyte differentiation. When treated with phorbol esters such as phorbol-myristate acetate (PMA), THP1 cells can differentiate into macrophages. Our initial approach utilizes RNA-seq technologies. This approach is highly appealing as it is cost-effective, and one can make use of publicly available datasets. Our second, more comprehensive approach is the screening method, which impartially evaluates all lncRNAs and their involvement in monocyte differentiation by CRISPR interference (CRISPRi) transcriptional silencing. For each candidate, we examined changes in open chromatin as indicated by previously published ATAC-seq before and after PMA treatment, as well as the presence of H3K27ac active chromatin marks near the putative TSS (13). We examine the advantages and disadvantages of these methodologies and the emergence of novel research and technologies aiding in addressing these challenges.

## Results

### Approaches to identify functional lncRNAs in human monocyte-to-macrophage differentiation

Here, we present two functional pipelines designed to identify lncRNAs that regulate monocyte-to-macrophage differentiation as outlined in Fig. 1A. The first approach utilizes RNA-seq to identify the most upregulated lncRNAs following differentiation and functional characterization to determine if these lncRNAs also play a role in the differentiation process. The second approach involves unbiased high throughput CRISPRi screening targeting all lncRNAs expressed in THP1 monocytic cells to determine which are involved in the differentiation process.

**Figure 1:**
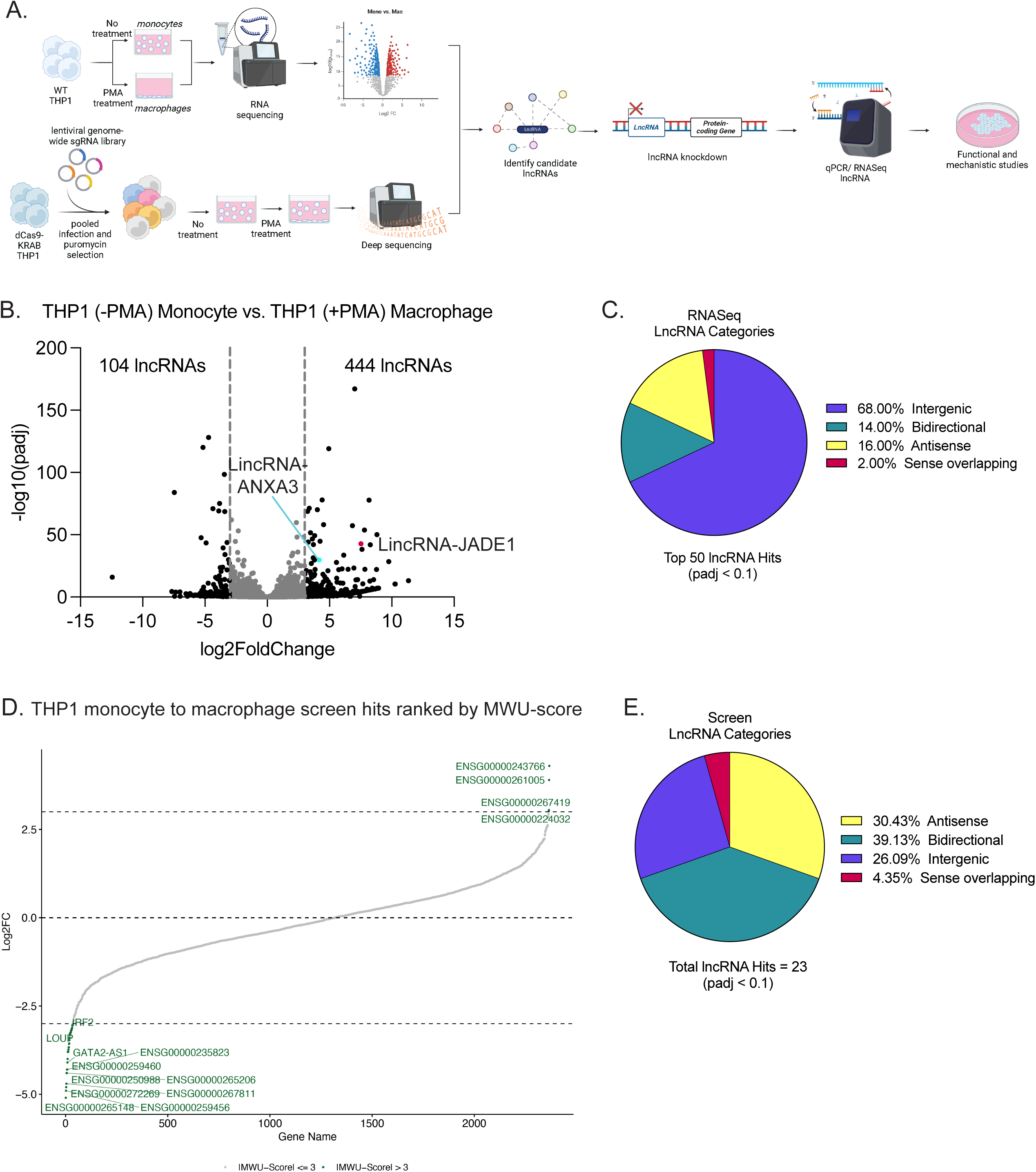
Approaches to identify functional lncRNAs in human monocyte to macrophage differentiation. **A. RNA sequencing and monocyte differentiation screen identify lncRNAs involved in monocyte-to-macrophage differentiation.** Wildtype THP1 cells were separated into two different groups: a no-treatment monocyte group and a PMA-treated macrophage group. RNA sequencing was performed on both groups, followed by DESeq2 analysis. dCas9-KRAB expressing THP1 cells were infected with a pooled sgRNA library and selected using puromycin. Three days post-infection, cells were harvested to collect the no-treatment time point. Cells were selected for 7 days and collected again 24hrs after PMA treatment. SgRNAs from each time point were PCR amplified and sequenced. Candidate lncRNAs from both approaches converged into one pipeline consisting of lncRNA knockdown experiments followed by qPCR/ RNA sequencing and functional and mechanistic studies. **B. THP1 monocyte vs. macrophage RNA sequencing analysis.** DESeq2 was used to establish log2foldchange of lncRNAs between no-treatment and PMA-treated groups to identify upregulated and downregulated lncRNAs. LFCs of −3 and 3 were considered significant. The magenta circle is *LincRNA-JADE1* and the blue circle is *LincRNA-ANXA3*. **C. Top 50 upregulated lncRNA categories.** The top 50 upregulated lncRNAs were categorized based on lncRNA type using UCSC Genome Browser (Human hg38). **D. THP1 monocyte to macrophage screen analysis.** DESeq2 was used to establish log2foldchange (L2FC) of sgRNAs between no treatment and PMA conditions. L2FC for sets of sgRNAs targeting each gene were compared to L2FC of all negative controls by Mann-Whitney U (MWU) test. MWU scores of −3 and 3 are considered significant. **E. Significant lncRNA screen hits categories.** The top lncRNA hits according to padj < 0.1 were categorized based on lncRNA type using UCSC Genome Browser (Human hg38).

### RNA-seq approach

For the first approach, THP1 cells’ RNA was sequenced as monocytes or macrophages (following treatment with 100nM PMA for 24 h). DESeq2 analysis comparing monocytes to macrophages identified 548 lncRNAs differentially expressed following differentiation (Fig.1B). 444 lncRNAs were increased in expression while 104 were reduced using a log2 foldchange (LFC) cutoff greater than 3 or less than −3 (Fig. 1B).

We established the following criteria to determine which candidates were worth pursuing further.

➢ Intergenic lncRNAs: Intergenic lncRNAs are characterized by having distinct promoters and being situated at least 1kb apart from adjacent protein-coding genes. We can study their functions independent of neighboring protein-coding genes. This enables us to utilize diverse molecular methods, including CRISPRi to determine function.
➢ Interesting Protein-Coding Gene Neighbors: A lncRNA with a neighbor shown or implicated to be involved in cell differentiation is interesting as it could indicate *cis* regulation between the lncRNA and neighboring protein. There are many examples of lncRNAs that regulate their protein-coding gene neighbors (*cis* regulation), such as *LincRNA-Cox2*, *UMLILO*, and *LOUP* (14, 15, 16).
➢ LncRNA Conservation: While lncRNAs generally exhibit poor conservation across species at a sequence level, we can evaluate their conservation by considering synteny (their position relative to conserved neighboring protein-coding genes) and the conservation of expression patterns.

For these reasons, we prioritized pursuing *LincJADE1* and *LincANXA3* from our RNA-seq studies. Of the top 50 lncRNAs, 34 lncRNAs were intergenic (defined as having a promoter at least 1kb away from a neighboring gene). Of these intergenic lncRNAs, 16 neighbor a protein-coding gene known or implicated in cell differentiation. It is possible that the other 18 intergenic lncRNAs neighbor a protein-coding gene yet to be identified as being involved in cell differentiation or that these lncRNAs function *in trans* to regulate the differentiation process (Fig. 1C).

### High throughput screening approach

We have previously described our CRISPRi screen to identify lncRNA regulators of monocyte-to-macrophage differentiation (8). In short, THP1 cells were infected with a pooled lentivirus and selected with puromycin for 7 days. Cells were treated three times over 11 days with 2nM PMA, allowing 50% of cells to differentiate (adhere to the plate) (8). This ensured the discovery of both positive and negative regulators of differentiation. 38 lncRNAs were identified with a Mann-Whitney U score greater than 3 or less than −3. 23 lncRNAs were found to have a p-adjusted value cutoff of less than 0.1 (Fig. 1D). Of our 23 statistically significant lncRNA hits, 9 were bidirectional; these lncRNAs shared a promoter and were transcribed from the opposite strand of their protein-coding gene neighbor. The second most prominent lncRNA category identified was antisense lncRNAs (Fig. 1E). Since our sgRNA library included lncRNAs regardless of genomic location, it is unsurprising that many of our hits are near or overlapping their protein-coding gene neighbor. We chose to pursue *GATA2-AS1* and *PPP2R5C-AS1* further because they neighbor protein-coding genes known to be involved in cell differentiation (17, 18). Interestingly, many of these bidirectional and antisense lncRNAs have protein-coding gene neighbors that have not previously been shown to play a role in monocyte differentiation, potentially making them novel coding regulators in this cellular process.

There were 7 lncRNA hits identified in both the RNA-seq and screening approaches (*HOTTIP, OLMALINC, ENSG00000264772, GATA2-AS1, LOUP, MIR17HG,* and *NUP153-AS1*), demonstrating that both of these approaches served to identify functional lncRNAs involved in monocyte differentiation. Of these hits, *OLMALINC* and *LOUP* are the only intergenic lncRNAs, but *LOUP* is the only lncRNA to neighbor a gene known to be involved in immune cell differentiation (*SPI1*) (8, 16).

### *LincRNA-JADE1* acts as a positive regulator of monocyte-to-macrophage differentiation

*LincRNA-JADE1* (*ENSG00000248187*) was identified as a highly inducible lncRNA following differentiation in our RNA-seq analysis and was selected for functional follow-up as it hit a number of our key criteria for selecting a hit: it is intergenic, neighbors an interesting protein-coding gene with possible roles in the differentiation process and shows synteny with mouse indicating possible conservation across species. To better understand the regulation at this locus and its possible mechanisms of action, we first examined the epigenetic landscape of the *LincJADE1* locus during differentiation (PMA treatment) (15). Epigenetic signatures, such as those detected through ATAC-seq, indicated a progressive expansion of chromatin accessibility and transcription activity at the transcription start site (TSS) of *LincJADE1* during differentiation (Fig. 2A). Furthermore, the active chromatin mark H3K27ac on the *LincJADE1* promoter suggests the potential for transcription factors to engage and trigger transcription at this site (Fig. 2A). Since some lncRNAs are multifunctional loci with the potential to encode short peptides from short open reading frames (sORFs) we also examined existing Ribo-Seq datasets from THP1 cells (8) and found that *LincJADE1* harbors an sORF with translation potential (Fig. 2A).

**Figure 2:**
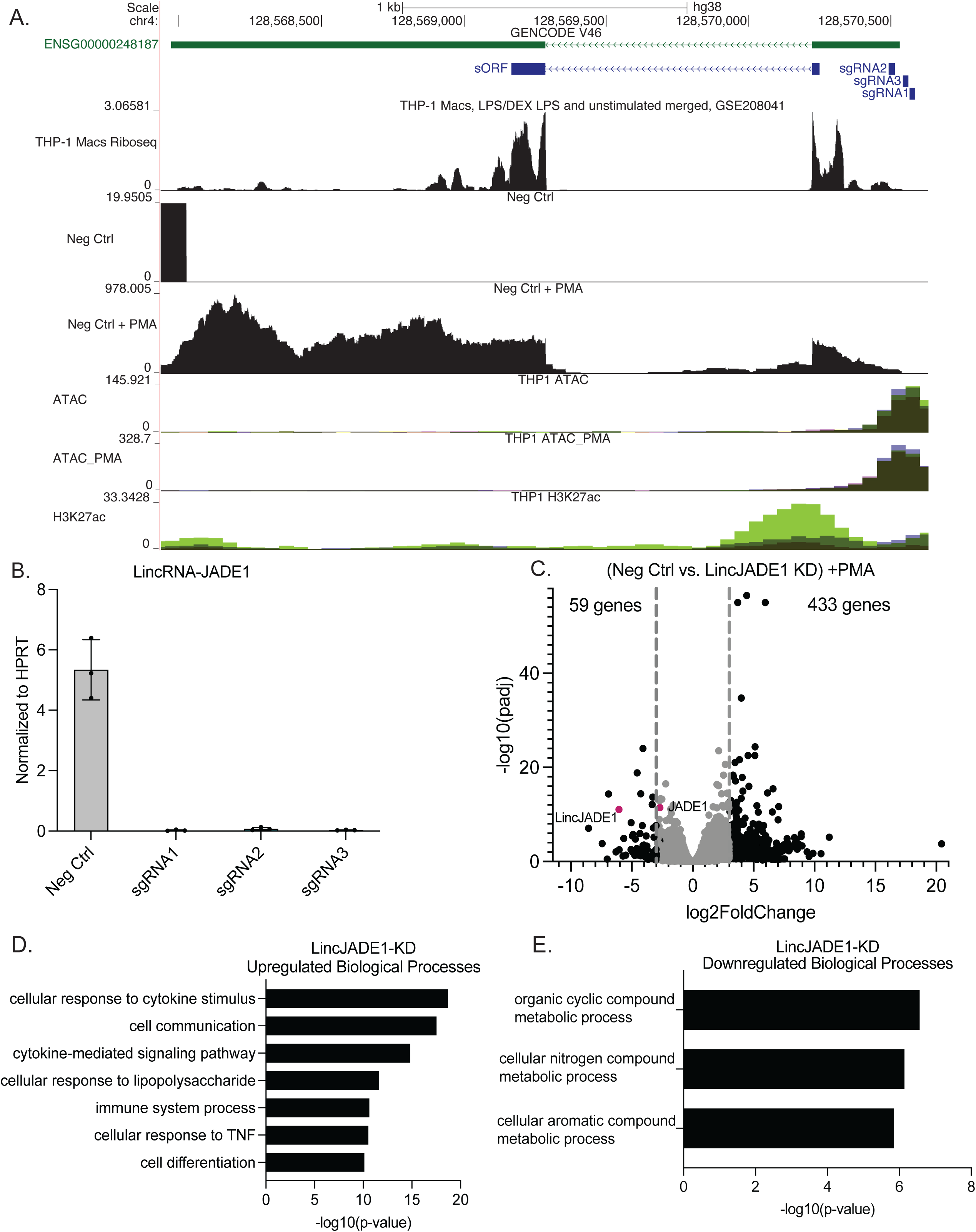
THP1 RNA sequencing identifies *LincRNA-JADE1* as a positive regulator of monocyte-to-macrophage differentiation. **A. SgRNAs targeting *LincJADE1***. The browser track displays the sORF present at the lncRNA locus and the 3 sgRNAs (sgRNA1, sgRNA2, sgRNA3) designed to knock down *LincJADE1* (*ENSG00000248187*). Browser tracks also display ATAC peaks in THP1 cells before and after PMA treatment, H3K27ac peaks, and Ribo-seq peaks in THP1 macrophages (THP-1 Macs GSE208041). Different colored peaks of ATAC-seq tracks represent different replicates. Numbers on the Y-axis represent raw read counts. **B. CRISPRi knockdown of *lincJADE1* in THP1 cells.** Three sgRNAs (sgRNA1, 2, 3) were designed to target *lincJADE1*. qPCR of *LincJADE1* across three technical replicates shows a statistically significant knockdown of *LincJADE1* by all three sgRNAs vs. a non-targeting sgRNA (Neg Ctrl). Values are normalized to HPRT, and error bars represent standard deviation. **C. Negative control vs. *LincJADE1*-KD RNA sequencing analysis.** DESeq2 was used to establish log2foldchange of genes between negative control and *LincJADE1* knockdown groups to identify upregulated and downregulated genes after PMA treatment. LFCs of −3 and 3 were considered significant. **D. Enrichment analysis of upregulated genes in *LincJADE1*-KD THP1 cells after PMA treatment.** Enriched GO terms of upregulated genes (LFC > 3) after knockdown of *LincJADE1.* **E. Enrichment analysis of downregulated genes in *LincJADE1*-KD THP1 cells after PMA treatment.** Top 3 enriched GO terms of downregulated genes (LFC < −2) after knockdown of *LincJADE1*.

To further test *LincJADE1*’s potential role as a positive regulator of monocyte differentiation, we designed 3 sgRNAs to knock it down using our CRISPRi THP1 cell line. QPCR showed more than 95% knockdown of the *LincJADE1* transcript (Fig. 2B). Next we performed RNA-seq to determine what genes and pathways are impacted during differentiation when *LincJADE1* was knocked down. We identified 491 differentially expressed genes (DEGs) with LFC cutoffs of −3 and 3 in the *LincJADE1* deficient cells (Fig. 2C). As expected, *LincJADE1* was one of the most significantly knocked down genes and, interestingly its neighboring protein *JADE1,* which is 235kb away, was also downregulated with an LFC of −2.7, hinting at a *cis-regulatory* relationship as well as a potentially undiscovered role for *JADE1* in monocyte differentiation.

Next, we performed a Gene Ontology (GO) analysis of the DEGs to determine which biological pathways *LincJADE1* might play a role in. The most significant upregulated genes (LFC > 3) were shown to be involved in cellular response to cytokine stimulus, cell communication, cytokine-mediated pathways, cellular response to lipopolysaccharide (LPS), immune system process, cellular response to tumor necrosis factor (TNF), and cell differentiation (Fig. 2D).

Altogether, these results show that *LincRNA-JADE1* is an intergenic lncRNA that is epigenetically primed upon PMA treatment to play a role in cell differentiation and the immune response.

### *LincRNA-ANXA3* is a positive regulator of monocyte-to-macrophage differentiation

Another promising novel lncRNA candidate identified from our THP1 RNA-seq was *LincANXA3* (*Linc01094*). *LincANXA3* is an intergenic lncRNA 86kb away from its nearest protein-coding gene neighbor, *ANXA3*. To enhance our understanding of the regulation at this locus and its potential mechanisms, we initially investigated the epigenetic landscape of the *LincANXA3* locus. ATAC-seq showed that differentiation increasingly primes the *LincANXA3* TSS for transcription activity. Additionally, the presence of H3K27ac marks across the locus implies the potential for transcription factors to interact with the TSS and other enhancer regions within the locus (Fig. 3A).

**Figure 3:**
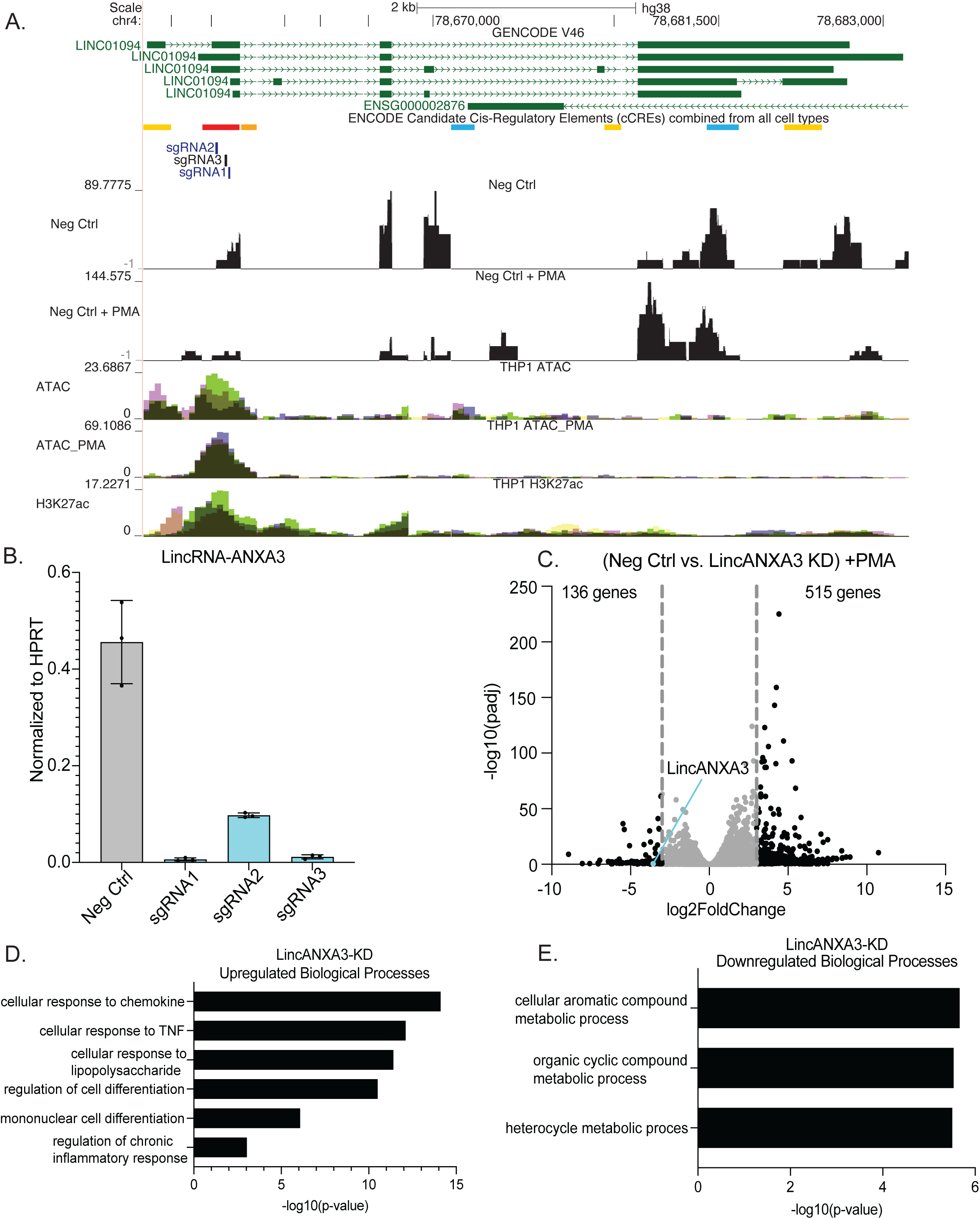
THP1 RNA sequencing identifies *LincRNA-ANXA3* as a positive regulator of monocyte-to-macrophage differentiation. **A. SgRNAs targeting *lincANXA3.*** The browser track displays 3 sgRNAs (sgRNA1, sgRNA2, sgRNA3) designed to knock down *LincANXA3* (*Linc01094*). ENCODE candidate cis-regulatory elements (cCREs) track displays promoter and enhancer-like signatures. Red bars represent promoter-like signatures, orange and yellow bars represent proximal and enhancer-like signatures, and blue bars represent CTCF binding signatures. Browser tracks also display ATAC peaks in THP1 cells before and after PMA treatment and H3K27ac peaks. Different colored peaks of ATAC-seq tracks represent different replicates. Numbers on the Y-axis represent raw read counts. **B. CRISPRi knockdown of *LincANXA3* in THP1 cells.** Three sgRNAs (sgRNA1, 2, 3) were designed to target *LincANXA3*. qPCR on *lincANXA3* across three technical replicates shows a statistically significant knockdown of *LincANXA3* by all three sgRNAs vs. a non-targeting sgRNA (Neg Ctrl). Values are normalized to HPRT, and error bars represent standard deviation. **C. Negative control vs. *LincANXA3*-KD RNA sequencing analysis.** DESeq2 was used to establish log2foldchange of genes between negative control and *LincANXA3* knockdown groups to identify upregulated and downregulated genes after PMA treatment. LFCs of −3 and 3 were considered significant. **D. Enrichment analysis of upregulated genes in *LincANXA3*-KD THP1 cells after PMA treatment.** Enriched GO terms of upregulated genes (LFC > 3) after knockdown of *LincANXA3.* **E. Enrichment analysis of downregulated genes in *LincANXA3*-KD THP1 cells after PMA treatment.** Top 3 enriched GO terms of downregulated genes (LFC < −2) after knockdown of *LincANXA3*.

Following our pipeline, we designed 3 sgRNAs to target *LincANXA3* using CRISPRi. According to Gencode, there are 5 different isoforms of this lncRNA. However, 4 of the 5 utilize the same TSS, with the first exon possessing H3K4me3 promoter-like signatures, in addition to our RNA-seq data indicating that this is the bonafide start site. Therefore, we designed our sgRNAs for this region. Using qPCR we found that we achieved 90% knockdown of the *LincANXA3* transcript (Fig. 3B). After validating *LincANXA3* knockdown, we performed RNA-seq. We identified 651 DEGs with LFC cutoffs of −3 and 3 (Fig. 3C). As expected, we saw *LincANXA3* downregulated, while its neighbor *ANXA3* did not change significantly with an LFC of 1.0 and an adjusted p-value greater than 0.1.

To determine whether *LincANXA3* could play a role in monocyte differentiation signaling circuits, we input DEGs into GO. The most significant upregulated genes (LFC < 3) were shown to be involved in cellular response to chemokine, cellular response to TNF, cellular response to LPS, regulation of cell differentiation, mononuclear cell differentiation, and regulation of chronic inflammatory response (Fig. 3D). These results show that *LincANXA3* is an intergenic lncRNA that is primed upon PMA treatment to enhance cell differentiation and the immune response.

Since its neighboring gene is not impacted when *LincANXA3* is removed suggests this gene functions to regulate differentiation in *trans* through mechanisms that still need to be determined.

### *GATA2-AS1* is a positive regulator of monocyte differentiation

*GATA2-AS1* was hit number 10 from our THP1 differentiation screen with an adjusted p-value less than 0.1. *GATA2-AS1* is an antisense lncRNA, meaning that it is transcribed on the opposite strand of its nearest-protein coding gene neighbor, *GATA2*. There are two isoforms of *GATA2-AS1*, one shorter isoform with 3 exons and a longer isoform with 2 exons. From ATAC-seq data, there does not appear to be a large difference in the epigenetic landscape between THP1 monocytes and THP1 macrophages (PMA treated), suggesting that this gene is transcriptionally active prior to differentiation. H3K27ac signals at the TSS of *GATA2-AS1* indicate this is also the case with strong transcription factor peaks in monocytes. However, due to the proximity of the lncRNA and *GATA2* protein coding gene it is difficult to determine if these marks relate to the lncRNA or the protein or both (Fig. 4A).

**Figure 4:**
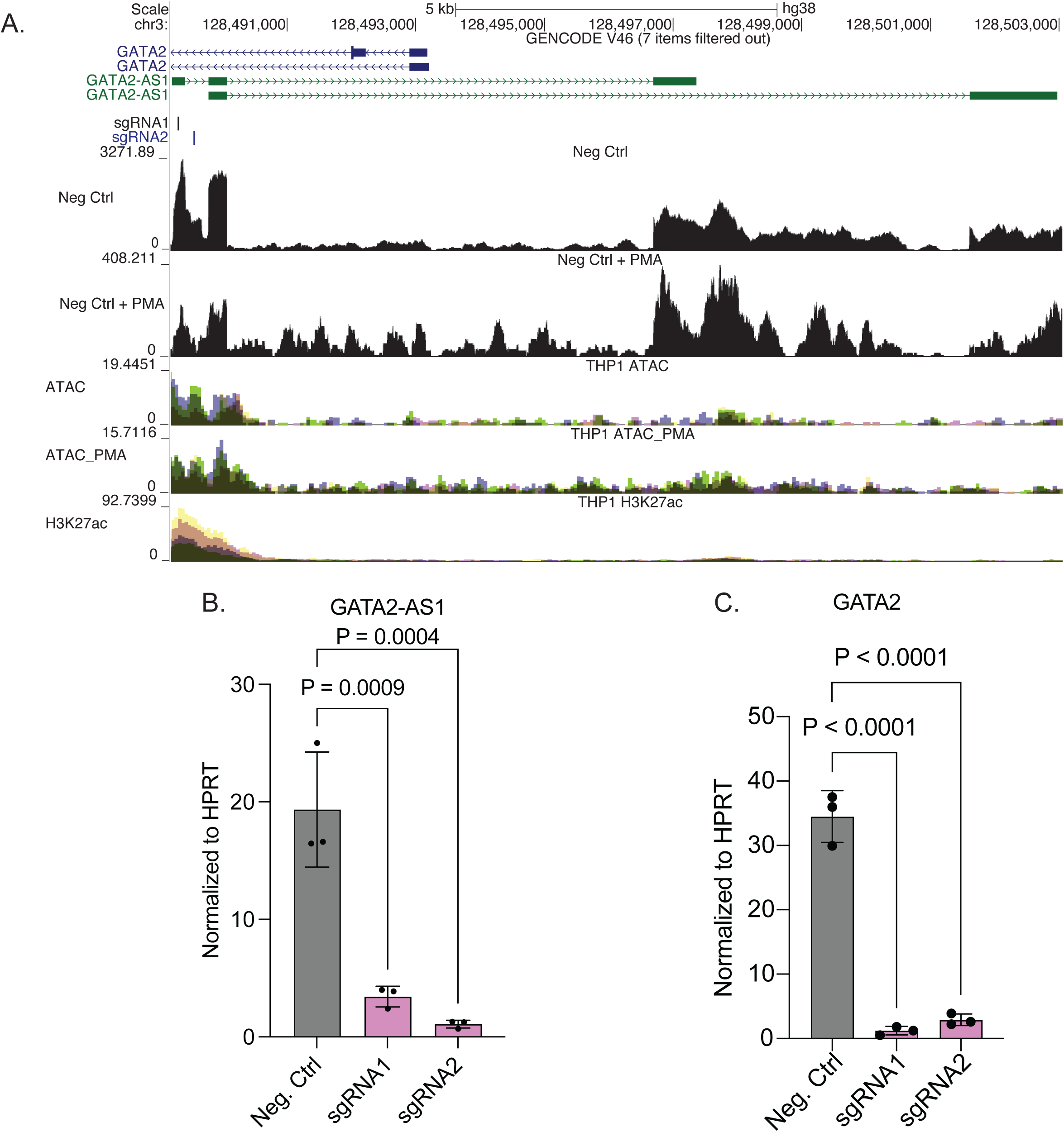
THP1 monocyte-to-macrophage screen identifies *GATA2AS1* as a positive regulator of monocyte differentiation. **A. SgRNAs targeting *GATA2AS1.*** The browser track displays the top 2 performing sgRNAs (sgRNA1 and sgRNA2) designed to knock down *GATA2AS1*. Browser tracks also display ATAC peaks in THP1 cells before and after PMA treatment and H3K27ac peaks. Different colored peaks of ATAC-seq tracks represent different replicates. Numbers on the Y-axis represent raw read counts. **B. CRISPRi knockdown of *GATA2AS1* in THP1 cells.** qPCR measurement of *GATA2AS1* across three technical replicates shows a statistically significant knockdown of *GATA2AS1* by all 2 sgRNAs vs. 1 non-targeting control (Neg. Ctrl.). Values are normalized to HPRT, and error bars represent standard deviation.

We chose the top 2 performing guides from the screen according to their adjusted p-values and cloned them to perform further knockdown analyses. Using qPCR we found that we were able to successfully knockdown 80% of the *GATA2-AS1* transcript (Fig. 4B). Since *GATA2-AS1*’s neighbor, *GATA2,* is already to be known to play a role in hematopoietic development and differentiation, we thought that perhaps the lncRNA could be part of the protein’s biological pathway because of their proximity and coexpression (17, 19). QPCR revealed over 90% knockdown of *GATA2* transcript, showing that it was also affected by CRISPRi knockdown. It is yet to be determined if this lncRNA is a hit because of the lncRNA itself or because CRISPRi heterochromatin silencing also silenced *GATA2,* and this is what made it a significant hit. Nonetheless, *GATA2* has yet to be shown to play a role in monocyte differentiation, potentially indicating a novel role for the protein also in this pathway.

### *PPP2R5C-AS1* is a positive regulator of monocyte differentiation

*PPP2R5C-AS1* (*ENSG00000259088*) was hit number 15 in our THP1 differentiation screen, with an adjusted p-value less than 0.1. This lncRNA is an antisense intronic lncRNA transcribed within one of *PPP2R5C*’s introns. ATAC-seq revealed that *PPP2R5C-AS1* possesses its own transcription start site (TSS), situated at a minimum distance of 48kb from the TSS of its neighboring protein-coding gene, *PPP2R5C*. There is a slight increase in ATAC-seq peaks following PMA treatment, indicating a possible change in the epigenetic landscape after differentiation (Fig. 5A). Interestingly, the H3K27ac signals at the TSS of *PPP2R5C-AS1* appear to be higher compared to those at the TSS of *PPP2R5C*. With the observed elevated RNA transcription at the TSS of *PPP2R5C-AS1*, it remains uncertain what function the transcript of this lncRNA serves in monocyte differentiation or whether the locus of the lncRNA acts as an enhancer to stimulate the activation of *PPP2R5C*.

**Figure 5:**
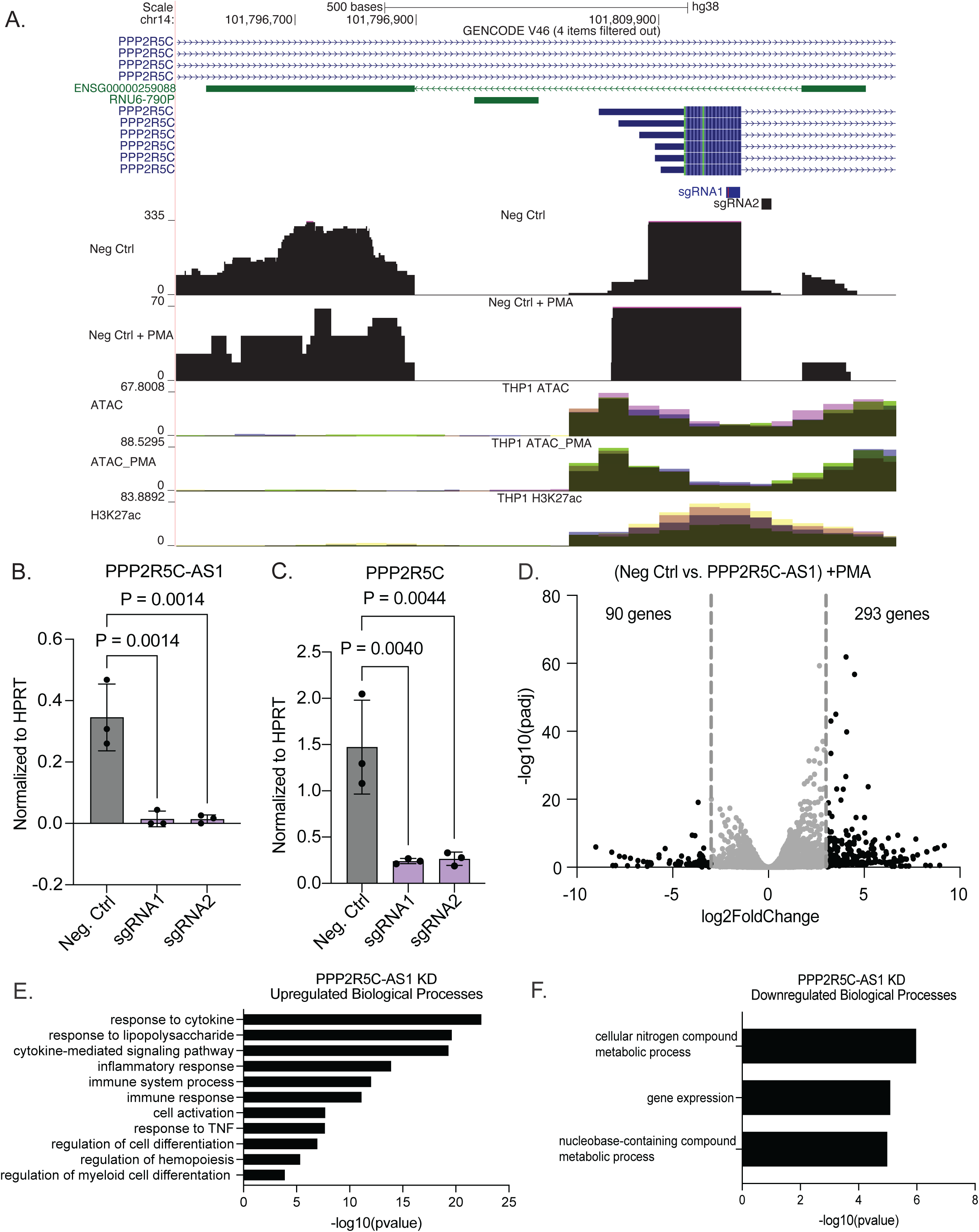
THP1 monocyte-to-macrophage screen identifies *PPP2R5C-AS1* as a positive regulator of monocyte differentiation. **A. SgRNAs targeting *PPP2R5C-AS1.*** The browser track displays the top 2 performing sgRNAs (sgRNA1 and sgRNA2) designed to knock down *PPP2R5C-AS1*. Browser tracks also display ATAC peaks in THP1 cells before and after PMA treatment and H3K27ac peaks. Different colored peaks of ATAC-seq tracks represent different replicates. Numbers on the Y-axis represent raw read counts. **B. CRISPRi knockdown of *PPP2R5C-AS1* in THP1 cells.** qPCR measurement of *PPP2R5C-AS1* across three technical replicates shows a statistically significant knockdown of *PPP2R5C-AS1* by all 2 sgRNAs vs. 1 non-targeting control (Neg. Ctrl.). Values are normalized to HPRT, and error bars represent standard deviation. **C. Negative control vs. *PPP2R5C-AS1* KD RNA sequencing analysis.** DESeq2 was used to establish log2foldchange of genes between negative control and *PPP2R5C-AS1* knockdown groups to identify upregulated and downregulated genes after PMA treatment. LFCs of −3 and 3 were considered significant. **D. Enrichment analysis of upregulated genes in *PPP2R5C-AS1* KD THP1 cells after PMA treatment.** Enriched GO terms of upregulated genes (LFC > 3) after knockdown of *PPP2R5C-AS1.* **E. Enrichment analysis of downregulated genes in *PPP2R5C-AS1* KD THP1 cells after PMA treatment.** Top 3 enriched GO terms of downregulated genes (LFC < −2.8) after knockdown of *PPP2R5C-AS1*.

We cloned the top 2 performing sgRNAs of *PPP2R5C-AS1*. Using qPCR, we found that there was almost 95% knockdown of *PPP2R5C-AS1* (Fig. 5B). Once we validated knockdown, we performed RNA-seq and identified 383 DEGs with a LFC cutoff of −3 and 3 (Fig. 5D). QPCR showed that *PPP2R5C* was significantly downregulated (Fig. 5C). However, it was not the most downregulated gene in our DESeq2 analysis. We input the upregulated genes with an LFC cutoff of greater than 3 into GO. The most significantly upregulated genes were shown to be involved in response to cytokine, response to LPS, cytokine-mediated pathway, inflammatory response, immune system response, cell activation, response to TNF, regulation of cell differentiation, regulation of hematopoiesis, and regulation of myeloid cell differentiation (Fig. 5D). Taken together, these results indicate that the *PPP2R5C-AS1* locus plays a role in monocyte differentiation. As with any antisense lncRNA hit, it is unknown whether the lncRNA transcript, protein-coding gene transcript, or a combination of both plays a crucial role in cell phenotype and makes it a significant hit. Further mechanistic work is required to confirm a novel role for *PPP2R5C* in monocyte differentiation.

## Discussion

The goal of this study is to compare two approaches to identify novel functional lncRNAs during monocyte-to-macrophage differentiation. THP1 monocytic cells can be differentiated using phorbol-12-myristate-13-acetate (PMA). This approach has been shown to result in comparable cytokine, metabolic, and differential gene expression to human peripheral blood mononuclear cells (PBMCs) (20). Numerous studies have been conducted to compare the timing and dosage of PMA’s transcriptomic effects during THP1 differentiation to establish a standardized protocol for researchers utilizing these cells. Although the optimal protocol remains undetermined across all experiments, these papers have effectively mapped out changes in the THP1 transcriptome induced by PMA-induced differentiation (20, 21). We have used this well-studied approach to differentiate THP1 cells and identify functional lncRNAs involved in monocyte differentiation. Our approach recapitulated many of the same findings previously reported in Lui et al’s paper, including DEGs and GO term biological pathways, further confirming the transcriptomic effects PMA differentiation has on THP1 cells (21).

There are over 20,000 lncRNAs identified in the human genome to date. This number increases with each new release of Gencode making the job of functionally characterizing these lncRNAs a challenge. Here we take two approaches, the RNA-seq approach and a CRISPRi based screening approach to identify viable candidate lncRNAs involved in macrophage differentiation. In the table below we outline the pros and cons to each of these approaches.

| **RNA-sequencing approach for identifying functional lncRNAs** |  |
| --- | --- |
| **<u>Pros</u>** | **<u>Cons</u>** |
| • Cheap and quick to perform<br>• Can utilize publically available data<br>• Can identify novel previously unannotated lncRNAs | • Makes assumptions that changes in expression are linked to regulation of a pathway<br>• High chance of following a false positive lead<br>• Mechanism of action can be difficult to determine |
| **High throughput screening approach to identify functional lncRNAs** |  |
| **<u>Pros</u>** | **<u>Cons</u>** |
| • Identifies hits based on function<br>• Produces high confidence hits | • Expensive, labor intensive and requires strong molecular biology |
|  | skills<br>• Slow can take 6-8 months<br>• Limited to a specific readout that works with pooled screening<br>• Targeting lncRNAs near coding genes is a challenge |

The RNA-seq approach to identifying functional lncRNAs is appealing as it is cost-effective and there are many public datasets that could be utilized to this end. The main hypothesis used in this approach is that if a lncRNA is induced by a given inflammatory stimulus then it suggests that the gene could be involved in the same pathway. This assumption carries risks because cellular signaling circuits are intricate, and transcript expression may not always correlate with cell phenotype. We have utilized this approach successfully in the past to begin to study *LincRNA-Cox2* and *GAPLINC* in the immune response (22, 23). Here we have successfully employed the approach to show that both *LincJADE1* and *LincANXA3* function within the monocyte differentiation process. Our results indicate that *LincJADE1* likely functions *in cis* to regulate its neighboring protein *JADE1*. In contrast, knockdown of *LincANXA3* also impacts differentiation pathways, but based on RNA-seq data, this lncRNA does not impact its neighbor *ANXA3* suggesting that it functions *in trans* to regulate gene expression. Further mechanistic work is required to determine how both these lncRNAs play roles in regulating the differentiation process.

The screening approach thoroughly tests all expressed lncRNAs and their role in monocyte differentiation unbiasedly. One of the caveats to our screen is that by using CRISPRi we may have disrupted not just lncRNA but also protein-coding gene transcription if they are close by in genomic space. Our CRISPRi approach allowed us to discover new regulators of monocyte differentiation among lncRNAs as well as potential novel functions for protein-coding genes. LncRNAs that share a promoter or overlap with protein-coding genes, like *GATA2-AS1* and *PPP2R5C-AS1,* are challenging to functionally characterize. Much more work is needed to tease apart these loci and determine what functions are being mediated by the lncRNA versus the protein or if they are both involved as a network. These two hits are interesting to us as it is possible that the important neighboring proteins *GATA2* and *PPP2R5C* also play novel roles in differentiation. Indeed recently *PPP2R5C* has been shown to be upregulated in acute myeloid leukemia which could be connected to its having a functional role in regulating the differentiation pathways (18). Follow up work to mechanistically tease apart the *GATA2-AS1* and *PPP2R5C-AS1* loci could involve targeting the neighboring proteins with active CRISPR to determine if this recapitulates the phenotype indicating that the protein is involved. Different approaches could be used to knockdown the lncRNA transcript or protein including siRNA or antisense oligo approaches. This would provide mechanistic insight into which components of these loci are contributing to the given phenotype.

The two approaches yielded common hits including *HOTTIP, OLMALINC, ENSG00000264772, GATA2-AS1, LOUP, MIR17HG,* and *NUP153-AS1.* We have recently published on the mechanistic role that *LOUP* plays in regulating macrophage differentiation and NFkB signaling through regulating its neighboring protein coding gene *SPI1* (8). This work highlights how multifunctional a given lncRNA loci can be. Therefore, it is important to try and determine if it is the RNA, the act of transcription itself or DNA elements or small proteins that are mediating the effects from the locus (5–7). The single lncRNA can function by one mechanism to regulate its neighbor *in cis* and it can move away from its site of transcription to regulate genes *in trans*. We have found this to be the case with *lincRNA-Cox2* that functions *in cis* to regulate its neighboring protein *PTGS2* through an enhancer RNA mechanism as well as functioning in *trans* to regulate a large number of immune genes (14, 22).

There are a number of technical challenges to the study of lncRNAs and their mechanisms of action. Here we propose two approaches to finding suitable candidates for future studies of lncRNAs involved in monocyte differentiation. The two approaches employed here primarily yielded distinct sets of lncRNA hits and our analysis revealed that our four selected candidate lncRNAs significantly influenced similar biological processes. These processes include cellular responses to cytokines/chemokines, inflammatory responses, and cell differentiation. Our lncRNA candidates may all regulate the same monocyte differentiation pathway, but their specific functions and mechanisms of action are yet to be determined. Both approaches can be applied equally to the study of lowly or highly expressed transcripts. RNA-seq lends itself to be applied in a variety of circumstances to identify genes involved in different biological processes. While screens can be more difficult to design outside of identifying viability genes, they provide more functional data. However, these pipelines can be easily adapted for use in any biological process, which will greatly aid in determining functional lncRNAs.

## Methods

### Cell lines

Wildtype (WT) THP1 cells were obtained from ATCC. All THP1 cell lines were cultured in RPMI 1640 supplemented with 10% low-endotoxin fetal bovine serum (ThermoFisher), 1X penicillin/streptomycin, and incubated at 37°C in 5% CO2. Cells were also treated with 100nM of PMA for 24 hr.

### Lentivirus production

All constructs were cotransfected into HEK293T cells with lentiviral packaging vectors psPAX (Addgene cat#12260) and pMD2.g (Addgene cat#12259) using Lipofectamine 3000 (ThermoFisher cat# L3000001) according to the manufacturer’s protocol. Viral supernatant was harvested 72h post-transfection.

### THP1-NFkB-EGFP-dCasKRAB

We constructed a GFP-based NF-κB reporter system by adding 5x NF-κB-binding motifs (GGGAATTTCC) upstream of the minimal CMV promoter-driven EGFP. THP1s were lentivirally infected and clonally selected for optimal reporter activity. Reporter cells were then lentivirally infected with the dCas9 construct that was constructed using Lenti-dCas9-KRAB-blast, addgene#89567. Cells were clonally selected for knockdown efficiency greater than 90%.

### THP1-NfKB-EGFP-dCASKRAB-sgRNA

NFkB-EGFP-CRISPRi-THP1 cells were lentivirally infected with sgRNAs. sgRNA constructs were made from a pSico lentiviral backbone driven by an EF1a promoter expressing T2A flanked genes: puromycin resistance and mCherry. sgRNAs were expressed from a mouse U6 promoter.Twenty-nucleotide forward/reverse gRNA oligonucleotides were annealed and cloned via the AarI site.

### Screening Protocol

#### sgRNA library design and cloning

10 sgRNAs were designed for each TSS of hg19 annotated lncRNAs expressed in THP1s at baseline and upon stimulation. The sgRNA library also included 700 non-targeting control sgRNAs, and sgRNAs targeting 50 protein coding genes as positive controls. The sgRNA library was designed and cloned as previously described (24).

### CRISPRi PMA Screen

THP1-NFkB-EGFP-CRISPRi-sgRNA were infected with the sgRNA library as previously described (8). Cell lines were infected, and the initial coverage after infection was ∼500 to 600×. Then, cells were expanded to >1,000× coverage. Triplicates were either left untreated or treated with 2 nM PMA on days 0, 8, and 9. Undifferentiated cells were collected on day 11, and sgRNAs were PCR amplified (8, 24).

### PMA Screen Analysis

SgRNAs were counted and passed to DESeq2 for analysis. Default normalization was performed, and log2foldchange (L2FC) was calculated for each sgRNA between the PMA and No Treatment conditions. L2FC for the set of sgRNAs targeting each gene were compared to L2FC of all negative controls by Mann-Whitney U (MWU) test. PMA replicate B was excluded from the analysis as it fell below 500X sgRNA coverage over the course of the experiment. Data are available at GSE247761 (8).

### RNA Isolation and RT-qPCR

Cells were homogenized in Tri-Reagent (Sigma Aldrich, T9424-200 mL). RNA was extracted with Direct-zol RNA miniprep plus RNA extraction Kit (Zymo, R2072). 1 ug of total RNA was reverse transcribed into cDNA (iScript cDNA synthesis kit, Bio-Rad cat# 1708840). cDNA was diluted 1:30 in qPCR experiments. Primers are outlined below

### Sequencing Data

RNA-seq was performed in wildtype THP1 cells (monocytes) and PMA-treated THP1 cells (macrophages). Data were originally reported in (23) and are available in GSE15057. Data pertaining to ATACSeq and ChIPSeq in THP1s were originally reported in (13) and are available at GEO: GSE96800 and SRA: PRJNA385337. Ribo-Seq data are from (24), with data available at GSE208041. For more information on how these data were processed please refer to the following reference (8). RNA-seq was performed on THP1-NFkB-EGFP-CRISPRi-sgRNA controls (nontargeting and anti-GFP) versus CRISPRi-lncRNA knockdown in monocytes (THP1s) and PMA-treated cells (macrophages). THP1 cells were stimulated with 100 nM PMA for 24 h. Total RNA (1 μg) was used to generate libraries using the Bio kit. 400ng of total RNA was sent out to Novogene for library preparation and sequencing using the Illumina NovoSeq 6000 as paired-end 150-bp reads.Transcripts were aligned to the human genome (assembly GRCh38/hg38). Data are available at GSE270346.

### Primers and Reagents

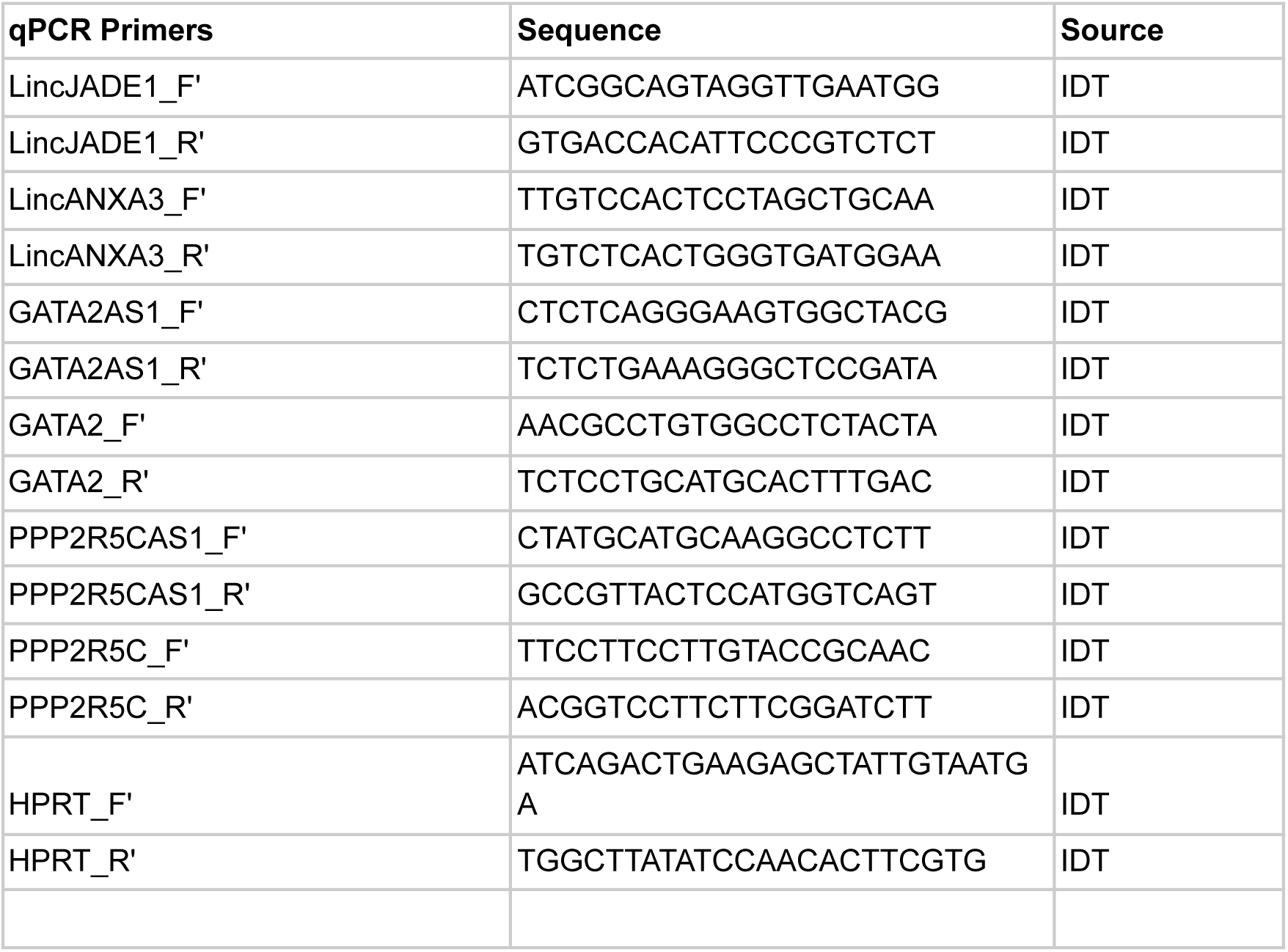
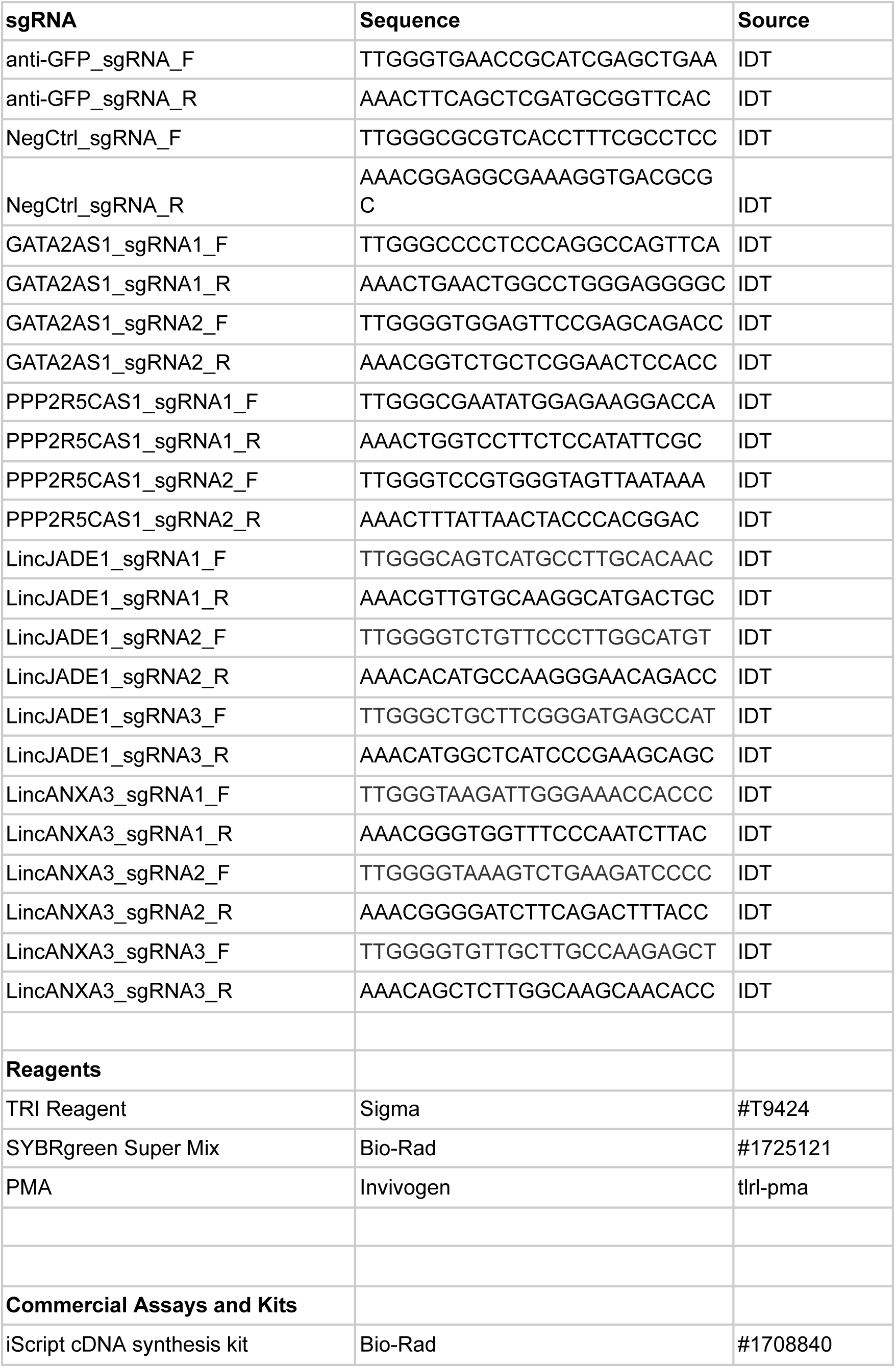
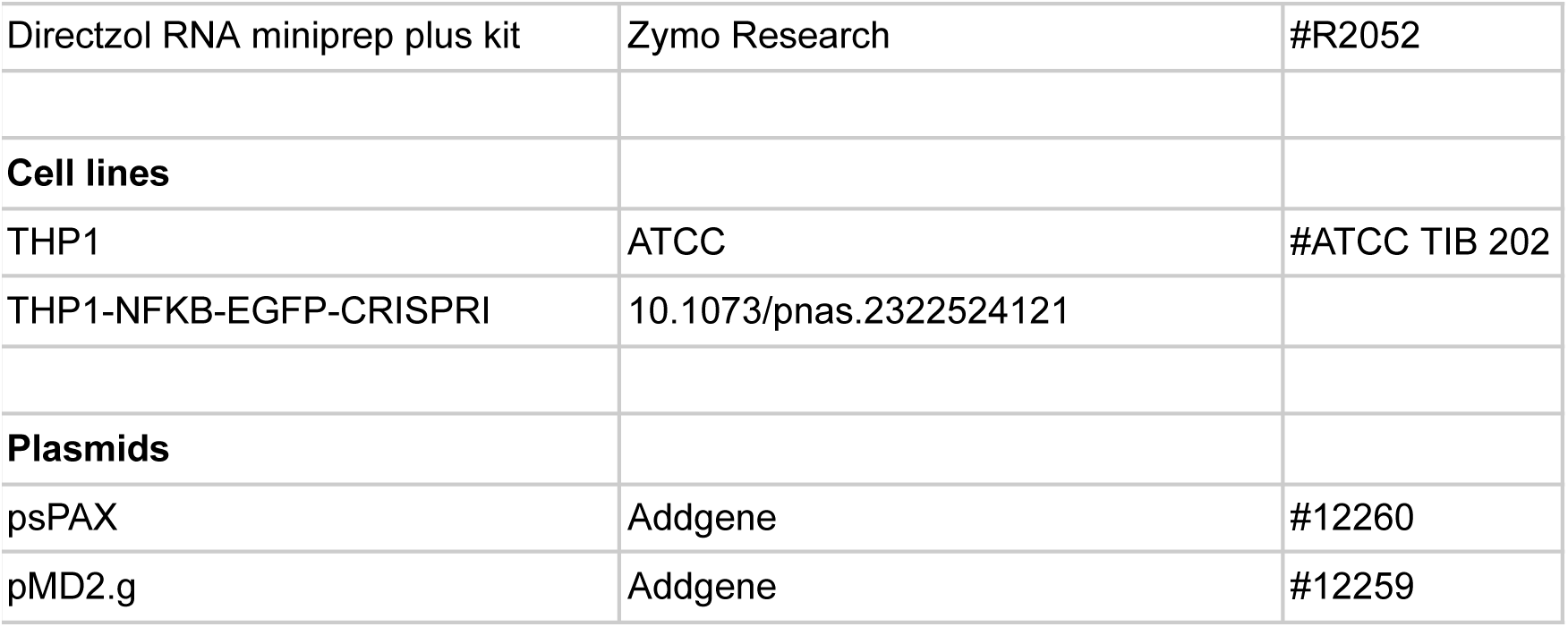

## Acknowledgements

S. Carpenter is supported by R35GM137801 from NIGMS, and C.M. is supported by a diversity supplement to R35GM137801 from NIGMS. E.M. is supported by F31AI179201 from NIAID.

## Author Contributions

C.M, S.Co, E.M and S.C designed the research. S.Co and S.C performed the CRISPR based screen and the initial RNA-sequencing. C.M performed the validation and downstream molecular mechanistic experiments of hits from the screens and RNA-seq approaches. E.M, S.Co and S.K performed the bioinformatic analysis to support the RNA-seq and screening studies. All authors contributed to the writing and editing of the manuscript.

## Competing interests

Carpenter is a paid consultant for NextRNA therapeutics.

